# Field synopsis and systematic meta-analyses of genetic association studies in isolated dystonia

**DOI:** 10.1101/306522

**Authors:** Olena Ohlei, Valerija Dobricic, Katja Lohmann, Christine Klein, Christina Lill, Lars Bertram

## Abstract

**Background and objectives:** Dystonia is a genetically complex disease with both monogenic and polygenic causes. For the latter, numerous genetic associations studies have been performed with largely inconsistent results. The aim of this study was to perform a field synopsis including systematic meta-analyses of genetic association studies in isolated dystonia

**Methods:** For the field synopsis we systematically screened and scrutinized the published literature using NCBI’s PubMed database. For genetic variants with sufficient information in at least two independent datasets, random-effects meta-analyses were performed, including meta-analyses stratified by ethnic descent and dystonia subtypes.

**Results:** A total of 3,575 articles were identified and scrutinized resulting in the inclusion of 42 independent publications allowing 134 meta-analyses on 45 variants across 17 genes. While our meta-analyses pinpointed several significant association signals with variants in *TOR1A, DRD1*, and *ARSG*, no single variant displayed compelling association with dystonia in the available data.

**Conclusions:** Our study provides an up-to-date summary of the status of dystonia genetic association studies. Additional large-scale studies are needed to better understand the genetic causes of isolated dystonia.

## Introduction

Genetically, dystonia is a complex disease with the parallel occurrence of forms with and without causative genetic factors. Over the past decades more than two dozen loci have been proposed as monogenic causes of dystonia[1–3] and 32 confirmed genes (28 with “DYT” labels)[4,5] have been included in the new list of isolated, combined, and complex hereditary dystonia proposed by the International Parkinson and Movement Disorder Society’s (MDS) Task Force on Genetic Nomenclature in Movement Disorders[5]. Substantially less is known about the genetic underpinnings of sporadic (i.e. non-monogenic) dystonia. Along these lines, only two GWAS on dystonia subtypes, i.e. musician’s dystonia[6] and cervical dystonia[7], have been published to date. The remainder of the existing genetics literature on sporadic dystonia is comprised of candidate gene association studies. These typically focused on genes known to cause familial dystonia as well as functionally founded candidates, such as genes involved in dopamine metabolism or the brain-derived neurotrophic factor (*BDNF*)[8]. To date, no systematic review covering all genetic polymorphisms investigated in dystonia has been published and only very few meta-analyses utilizing genetic association data exist[9–12]. As a result, it is becoming increasingly difficult to evaluate and interpret the genetics literature pertaining to sporadic dystonia. The aim of this work was to overcome this limitation and perform the first systematic synopsis - including meta-analyses of all available genetic association data - in the dystonia field. To this end, we carefully screened more than 3,500 articles and performed a total of 134 meta-analyses across 52 case-control datasets from 42 independent publications, thereby vastly increasing the number of meta-analyses currently available for dystonia[9–12]. The accrued literature database and meta-analysis results are made available in the supplement of this manuscript.

## Methods

### 1. Literature searches

#### 1.1. Systematic literature searches

Overall, our study followed the approach developed earlier by our group for systematic field synopses in Alzheimer’s disease[13] and Parkinson’s disease[14]. At the outset, this entailed a systematic literature search using NCBI’s “PubMed” database for papers published until January 23, 2018. In a first stage, we searched for keywords “(dystoni* OR (writer* AND cramp) OR graphospasm OR blepharospasm OR torticollis OR meige OR spasmodic OR dysphonia OR retrocollis OR antecollis OR laterocollis OR (musician* AND cramp) OR (occupation* AND cramp) OR (golf* AND cramp) OR yips) AND associat* AND gene*”. In the second stage, we separately searched for each gene identified in stage 1 and for 28 loci with a “DYT” label (Figure 1; DYT designations taken from refs. 5 and 15; see Supplementary Tables S1 and S2 for more details). Of note, prior to the revision of the dystonia classification and nomenclature in 2013, isolated dystonia (i.e. pure dystonia with or without dystonic tremor) was referred to as “primary” dystonia, a term mixing phenomenological and etiological features, whose use is no longer recommended[16]. As the clinical part of the definition of “primary” dystonia was identical to that of “isolated” dystonia, we exclusively use the new and unambiguous term “isolated” dystonia for all studies referred to in this article.

**Figure 1.**
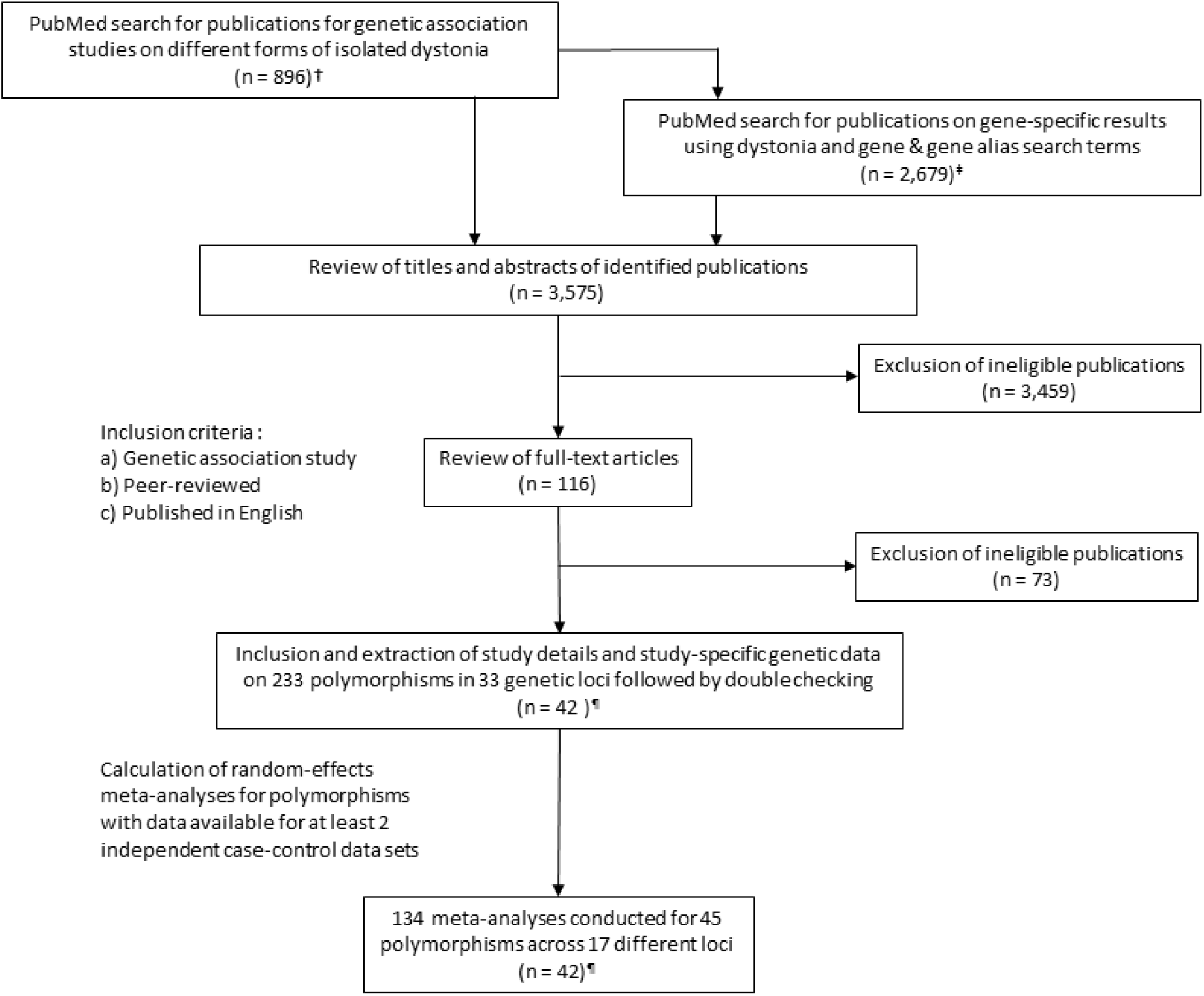
Flowchart of literature search, data extraction, and analysis strategies applied for the dystonia field synopsis. † Search for query terms “(dystoni* OR (writer* AND cramp) OR graphospasm OR blepharospasm OR torticollis OR meige OR spasmodic OR dysphonia OR retrocollis OR antecollis OR laterocollis OR (musician* AND cramp) OR (occupation* AND cramp) OR (golf* AND cramp) OR yips) AND associat* AND gene*”. ‡ Search for query terms outlined above and “AND ([gene name] OR [gene alias(es)])” using HGNC (HUGO Gene Nomenclature Committee at the European Bioinformatics Institute) nomenclature^27^. ¶ Only 45 out of 233 polymorphisms were investigated in at least two independent data sets. Furthermore, two studies were excluded due to sample overlap: 1. Djarmati et al, 2009 (ref. 19) fully excluded due to overlap with Lohman et al, 2012 (ref. 21), 2. Newman et al, 2012 (ref. 18) exclusion of data for polymorphisms overlapping with Newman et al, 2014 (ref. 20; see Table S2 for more details).

#### 1.2. Inclusion / exclusion criteria

We only considered publications representing association studies between biallelic genetic polymorphisms (minor allele frequency in included data sets or in the general population ≥0.01 based on ref. 17) and isolated dystonia phenotypes. Additional eligibility criteria were: publication in a peer-reviewed journal, publication in English, and analysis in at least 10 dystonia cases and 10 control subjects. We excluded studies assessing dystonia with a known etiology (previously known as “secondary” dystonia), studies using family-based designs, and studies on mitochondrial DNA. Furthermore, we excluded studies that contrasted their case groups against “database controls” and those with cases carrying a known dystonia mutation.

### 2. Database

From each eligible article, we extracted a range of informative variables into a Microsoft Excel database (full database in Supplementary Table S2). The initial data entry was performed by O.O. followed by double checking of all variant IDs and genotype distributions by V.D.

### 3. Statistical analyses

For all eligible datasets, we calculated odds ratios (OR) and corresponding 95% confidence intervals (CIs), assuming an additive genetic model. For datasets with allele frequency information only, we calculated the corresponding genotype distributions assuming Hardy-Weinberg equilibrium. For four variants, the same or largely overlapping data sets were published in separate articles[18–21]. In these instances, we only included the genotype distributions for the larger of the two datasets, i.e. refs. 20 and 21. For variants with independent genetic data in at least two non-overlapping datasets we calculated random effects summary ORs using PLINK 1.9[22,23]. Whenever sufficient data were available, meta-analyses were stratified by ethnic descent group in addition to calculating summary ORs across all ethnic groups combined. In addition to calculating meta-analyses across all forms of isolated dystonia, we divided our analyses into focal dystonia (i.e. cervical dystonia, blepharospasm, musician’s dystonia, writer’s cramp, and spasmodic dysphonia), segmental dystonia, and generalized dystonia. Whenever possible, subgroups were further stratified into groups of differing ethnic descent. Overall, this led to 134 meta-analyses of 45 variants across 17 genes. Pairwise linkage disequilibrium estimates across SNPs within the same ±1Mb interval in the 1000GP (phase 3) data[17] were calculated using PLINK 1.9 (Supplementary Table S3). This identified 32 independent (r^2^<0.3) variants; accordingly, the study-wide significance threshold was set to α = 0.00156 (=0.05/32). Heterogeneity across studies was assessed by calculating the I^2^ metric (I^2^ >75% indicates excess heterogeneity[14]).

## Results

### 1. Literature search

The systematic literature search (up-to-date until January 23rd, 2018) identified 3,575 potentially eligible articles (Figure 1). After systematic review of titles, abstracts and full text (as needed) versions of these papers we identified 42 eligible publications (describing association data in 52 independent datasets. Analyzed datasets originated from a total of 17 different countries spread across four continents. In total, 233 polymorphisms across 33 loci were investigated in these 42 publications. Most (n=40) used a candidate gene approach, while two papers were GWAS. The entire database of this field synopsis is provided in Supplementary Table 2.

### 2. Meta-analysis results

For the meta-analyses, we only considered variants with at least two independent non-overlapping datasets with available genotype or allele data. This included screening 557,621 SNPs from a musician’s dystonia GWAS dataset[6] for an overlap with the 233 polymorphisms identified in the literature screen. Overall, this procedure resulted in sufficient data for a total of 45 variants across 17 genetic loci (median sample size per data set: 458, range 88-5,385). Of these, 42 variants could be analysed after stratification in for datasets of Caucasian descent, and 6 after stratification for Asian descent. Only five variants in *TOR1A, DRD1* and *ARSG* showed nominally significant evidence for association: i.e. rs4532 (OR [95 % CI]: 1.37 [1.13-1.67], P=0.00153), rs35153737 (OR [95% CI]: 1.46 [1.14-1.88], P=0.00315), rs13283584 (OR [95% CI]: 1.32 [1.08-1.61], P=0.00584), rs7342975 (OR [95 % CI]: 2.00 [1.15-3.47], P=0.00572), rs9972951 (OR [95 % CI]: 2.16 [1.06-4.40], P=0.0342), but only variant rs4532 in *DRD1* survived correction for multiple testing. Meta-analyses in subgroups stratified by ethnic descent did not reveal any additional associations (Table 1).

**Table 1.** Results of random-effects meta-analyses based on an additive model in all types of isolated (a.k.a. “primary”) dystonia.

| Chr-position | Gene | SNP | Allele | Ethnicity ‡ | Cases vs Controls<br>(# data sets/ # publications) | OR (95%CI) | P-Value | I<sup>2</sup> |
| --- | --- | --- | --- | --- | --- | --- | --- | --- |
| 1:236885200 | MRT | rs1805087 | G vs A | C | 227 vs 1072 (2/2) | 0.93 (0.66-1.29) | 0.5852 | 0.00 |
| 2:178433097 | PRKRA | rs6739142 | G vs A | C | 357 vs 1200 (2/2) | 0.99 (0.73-1.36) | 0.9910 | 39.77 |
| 2:178449632 | PRKRA | rs1967327 | C vs G | C | 357 vs 1200 (2/2) | 1.10 (0.86-1.42) | 0.4392 | 0.00 |
| 2: 218326846 | PNKD | rs10932774 | A vs G | C | 357 vs 1200 (2/2) | 1.02 (0.84-1.24) | 0.8736 | 0.00 |
| 3:114171968 | DRD3 | rs6280 | C vs T | C | 188 vs 200 (2/2) | 0.82 (0.61-1.11) | 0.1979 | 0.00 |
| 5:175441837 | DRD1 | rs155417 | T vs C | C | 188 vs 200 (2/2) | 0.73 (0.31-1.74) | 0.4792 | 0.00 |
| **5:175443147** | **DRD1** | **rs4532** | **C vs T** | **C** | **315 vs 1172 (3/3)** | **1.37 (1.13-1.67)** | **0.0015** | **0.00** |
| 7:94635805 | SGCE | rs10235385 | C vs A | C | 357 vs 1200 (2/2) | 0.90 (0.69-1.16) | 0.4028 | 0.00 |
| 8:42838423 | THAP1 | rs111989331 | C vs T | All | 499 vs 360 (2/2) | 0.69 (0.31-1.54) | 0.3686 | 74.48 |
| 8:42842898 | THAP1 | rs71521601 | G vs A | C | 864 vs 510 (3/3) | 1.20 (0.66-2.21) | 0.5484 | 73.02 |
| 8:42843330-1 | THAP1 | rs370983900 & rs184497763 | AA vs CT | C | 3679 vs 2315 (7/7) | 1.07 (0.73-1.56) | 0.7431 | 40.61 |
| **9:129812583** | **TOR1A** | **rs13283584** | **T vs C** | **C** | **473 vs 749 (2/2)** | **1.32 (1.08-1.61)** | **0.0058** | **0.00** |
| 9:129813148 | TOR1A | rs3842225 | - vs C | All | 2287 vs 2503 (11/10) | 1.05 (0.91-1.21) | 0.4862 | 43.18 |
| 9:129813148 | TOR1A | rs3842225 | - vs C | C | 1835 vs 2083 (9/8) | 1.01 (0.86-1.18) | 0.9450 | 44.67 |
| 9:129813148 | TOR1A | rs3842225 | - vs C | A | 452 vs 420 (2/2) | 1.28 (0.99-0.65) | 0.0505 | 0.00 |
| **9:129813558** | **TOR1A** | **rs35153737** | **- vs C** | **A** | **452 vs 420 (2/2)** | **1.46 (1.14-1.88)** | **0.0031** | **0.00** |
| 9:129813781 | TOR1A | rs1182 | A vs C | All | 2213 vs 2526 (11/10) | 1.05 (0.85-1.30) | 0.6742 | 72.36 |
| 9:129813781 | TOR1A | rs1182 | A vs C | C | 1572 vs 2010 (8/7) | 1.01 (0.77-1.32) | 0.9607 | 79.00 |
| 9:129813781 | TOR1A | rs1182 | A vs C | A | 641 vs 516 (3/3) | 1.15 (0.90-1.45) | 0.2647 | 0.00 |
| 9:129816005 | TOR1A | rs11787741 | G vs A | C | 473 vs 749 (2/2) | 0.95 (0.77-1.17) | 0.6260 | 0.00 |
| 9:129818407 | TOR1A | rs13297609 | C vs G | All | 351 vs 428 (2/2) | 1.16 (0.89-1.52) | 0.2742 | 0.00 |
| 9:129818622 | TOR1A | rs1801968 | G vs C | All | 2075 vs 2699 (11/11) | 0.95 (0.78-1.16) | 0.6332 | 43.00 |
| 9:129818622 | TOR1A | rs1801968 | G vs C | C | 1428 vs 1925 (6/6) | 1.02 (0.87-1.20) | 0.7870 | 5.33 |
| 9:129818622 | TOR1A | rs1801968 | G vs C | A | 626 vs 494 (4/4) | 0.74 (0.44-1.26) | 0.2641 | 62.34 |
| 9:129822779 | TOR1A | rs2296793 | A vs G | All | 1692 vs 2277 (10/9) | 1.02 (0.87-1.18) | 0.8381 | 42.61 |
| 9:129822779 | TOR1A | rs2296793 | A vs G | C | 1454 vs 1946 (8/7) | 1.02 (0.86-1.21) | 0.8221 | 45.59 |
| 9:129822779 | TOR1A | rs2296793 | A vs G | A | 238 vs 331 (2/2) | 0.99 (0.60-1.64) | 0.9738 | 64.39 |
| 9:133636819 | DBH | rs2797849 | C vs G | C | 490 vs 3054 (2/2) | 1.09 (0.89-1.32) | 0.4079 | 38.64 |
| 9:133639458 | DBH | rs2797851 | A vs G | C | 490 vs 3054 (2/2) | 1.07 (0.87-1.34) | 0.5395 | 42.38 |
| 9:133657065 | DBH | rs1611131 | G vs A | C | 490 vs 3054 (2/2) | 1.12 (0.83-1.51) | 0.4528 | 69.85 |
| 9:133657152 | DBH | rs6271 | T vs C | C | 490 vs 3054 (2/2) | 2.97 (0.37-24.02) | 0.3002 | 96.73 |
| 9:133663599 | DBH | rs129886 | T vs C | C | 490 vs 3054 (2/2) | 1.01 (0.86-1.20) | 0.8704 | 0.00 |
| 10:28431147 | intergenic | rs1249277 | G vs C | C | 464 vs 5515 (2/2) | 0.77 (0.41-1.45) | 0.4215 | 89.34 |
| 11:27658369 | BDNF | rs6265 | T vs C | All | 1591 vs 2780 (7/7) | 1.09 (0.98-1.21) | 0.1165 | 0.00 |
| 11:27658369 | BDNF | rs6265 | T vs C | C | 1146 vs 2350 (5/5) | 1.05 (0.91-1.21) | 0.4936 | 7.52 |
| 11:27658369 | BDNF | rs6265 | T vs C | A | 445 vs 430 (2/2) | 1.19 (0.98-1.43) | 0.0759 | 0.00 |
| 11:48246304 | OR4X2 | rs67863238 | C vs G | C | 464 vs 5515 (2/2) | 2.17 (0.40-11.70) | 0.3482 | 96.50 |
| 11:113400106 | DRD2 | rs1800497 | A vs G | C | 222 vs 267 (3/3) | 0.89 (0.65-1.22) | 0.4758 | 0.00 |
| 11:113425564 | DRD2 | rs1079597 | T vs C | C | 188 vs 200 (2/2) | 1.00 (0.69-1.46) | 0.9913 | 0.00 |
| 11:113475529 | DRD2 | rs1799732 | - vs G | C | 188 vs 200 (2/2) | 0.74 (0.46-1.19) | 0.2064 | 0.00 |
| 13:101406511 | NALCN | rs1338041 | C vs A | All | 565 vs 5804 (3/3) | 0.90 (0.59-1.36) | 0.6014 | 89.79 |
| 13:101406511 | NALCN | rs1338041 | C vs A | C | 464 vs 5515 (2/2) | 0.75 (0.51-1.10) | 0.1445 | 83.78 |
| 13:101407929 | NALCN | rs9518385 | C vs A | C | 339 vs 6145 (2/2) | 0.73 (0.51-1.05) | 0.0901 | 76.89 |
| 13:101430922 | NALCN | rs61973742 | G vs A | All | 665 vs 5804 (3/3) | 1.38 (0.41-4.64) | 0.6001 | 96.35 |
| 13:101430922 | NALCN | rs61973742 | G vs A | C | 464 vs 5515 (2/2) | 1.85 (0.26-13.37) | 0.5245 | 97.41 |
| 14:54837362 | GCH1 | rs2145945 | A vs G | C | 357 vs 1200 (2/2) | 0.92 (0.66-1.27) | 0.6038 | 64.12 |
| 14:54839739 | GCH1 | rs10483639 | C vs G | C | 360 vs 2062 (2/2) | 1.09 (0.72-1.64) | 0.6990 | 76.69 |
| 14:54841286 | GCH1 | rs17739146 | G vs A | C | 357 vs 1200 (2/2) | 0.91 (0.66-1.27) | 0.5921 | 22.85 |
| 14:54888313 | GCH1 | rs998259 | T vs C | C | 357 vs 1200 (2/2) | 0.97 (0.74-1.27) | 0.7973 | 36.39 |
| 17:68368608 | ARSG | rs1558877 | T vs C | C | 357 vs 5272 (2/2) | 0.96 (0.78-1.18) | 0.6883 | 39.26 |
| 17:68368663 | ARSG | rs1558878 | C vs T | C | 357 vs 5272 (2/2) | 0.88 (0.75-1.02) | 0.0846 | 0.00 |
| 17:68386068 | ARSG | rs11655081 | C vs T | All | 2420 vs 3608 (9/1) | 1.33 (0.95-1.86) | 0.0989 | 77.20 |
| 17:68386068 | ARSG | rs11655081 | C vs T | C | 2243 vs 3409 (8/1) | 1.38 (0.92-2.08) | 0.1224 | 79.97 |
| **17:68395091** | **ARSG** | **rs7342975** | **G vs A** | **C** | **357 vs 5272 (2/2)** | **2.00 (1.15-3.47)** | **0.0057** | **0.00** |
| **17:68395135** | **ARSG** | **rs9972951** | **A vs G** | **C** | **357 vs 5272 (2/2)** | **2.16 (1.06-4.40)** | **0.0342** | **68.35** |
| 22:19950115 | COMT | rs5993883 | T vs G | C | 490 vs 3056 (2/2) | 0.90 (0.61-1.31) | 0.5671 | 83.57 |
| 22:19964609 | COMT | rs4646316 | T vs C | C | 490 vs 3056 (2/2) | 1.08 (0.92-1.27) | 0.3332 | 0.00 |
| 22:19968169 | COMT | rs9332377 | T vs C | C | 490 vs 3056 (2/2) | 1.11 (0.92-1.34) | 0.2678 | 0.00 |

‡C = Of Caucasian descent (i.e. more than 90% of individuals in a data set belongs to this descent group); A = Asian descent; All = All ethnic

Meta-analyses divided by diagnostic subgroup are summarized in Table 2 and yielded five nominally significant results, two of which were not observed in the analyses without diagnostic stratification, i.e. rs1801968 in *TOR1A* (OR [95% C.I.]: 3.10 [1.25-7.68], P=0.0142), associated with writer’s cramp, and rs11655081 in *ARSG* (OR [95% C.I.]: 4.42 [2.72-7.19], P=2.11e-09, respectively) associated with musician’s dystonia (Table 2). While the latter finding surpassed the threshold of study- and genome-wide significance, we note that this meta-analysis is only based on two independent datasets, both of which originate from the same (and only) paper on the topic[6]. The same variant was also assessed in independent datasets with other focal (i.e. cervical, blepharospasm; Table 2B, D) and segmental (Table 2G) forms of dystonia but did not show any evidence for association with these dystonia subtypes[6].

**Table 2.**
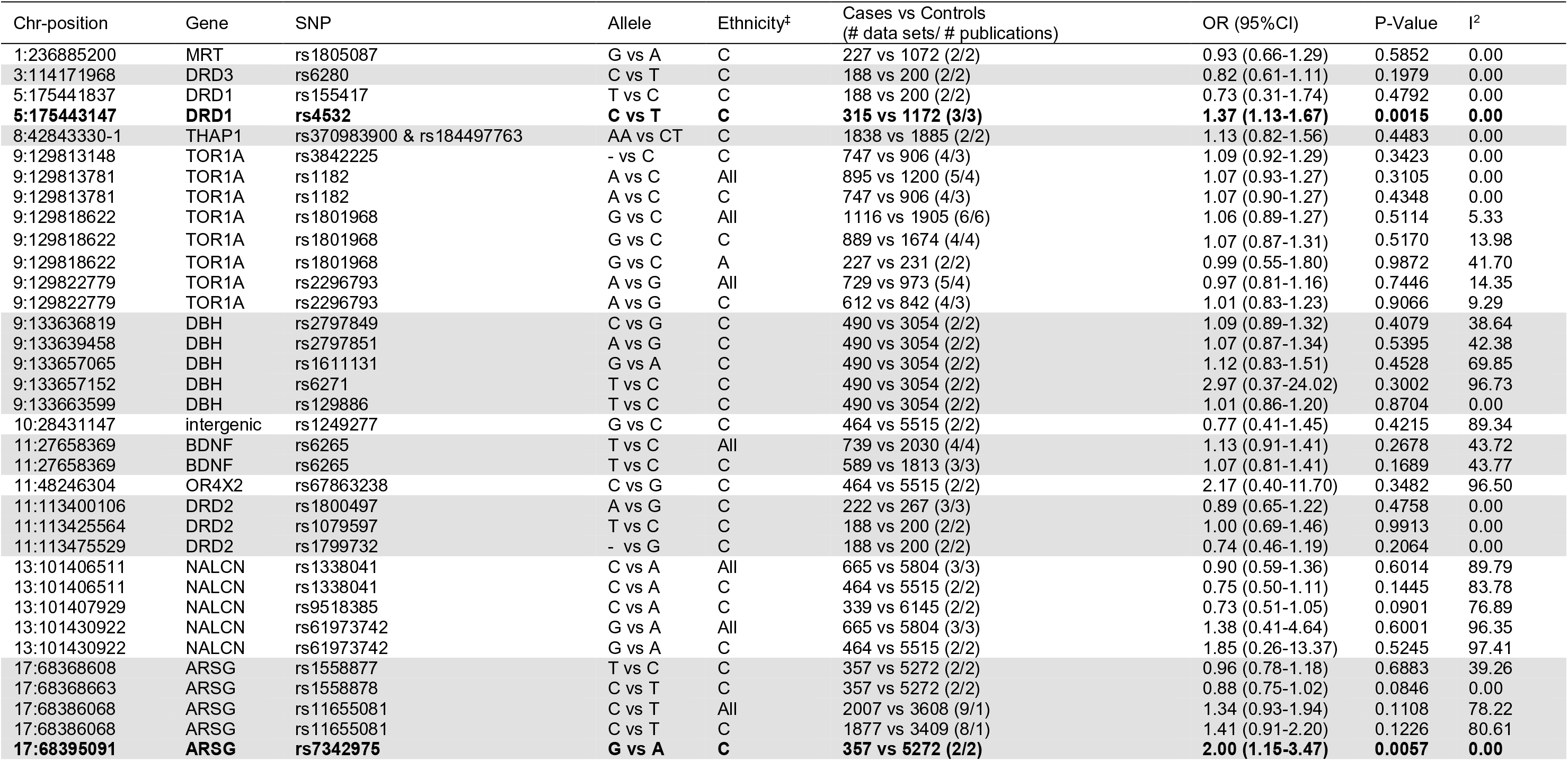
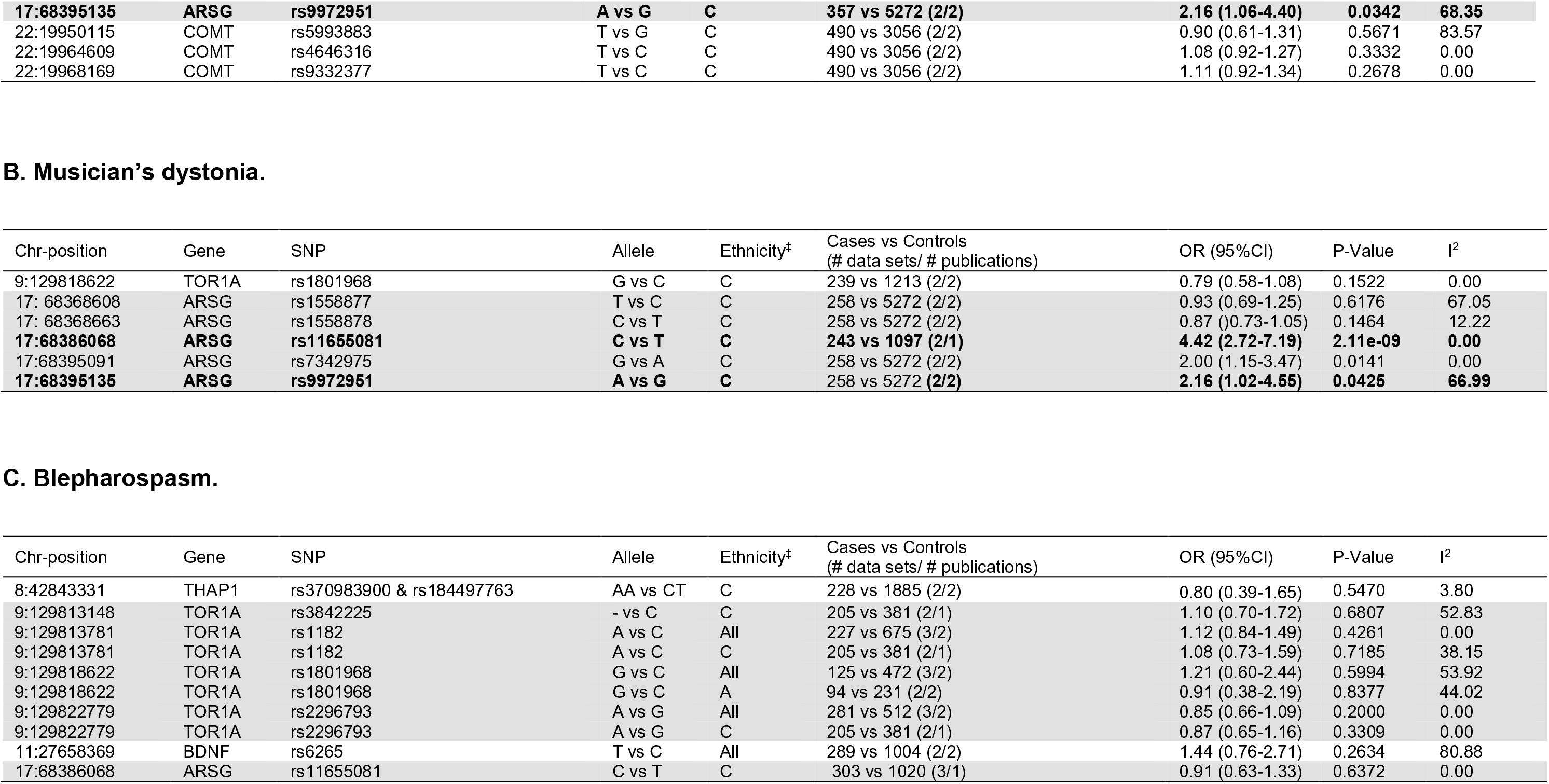
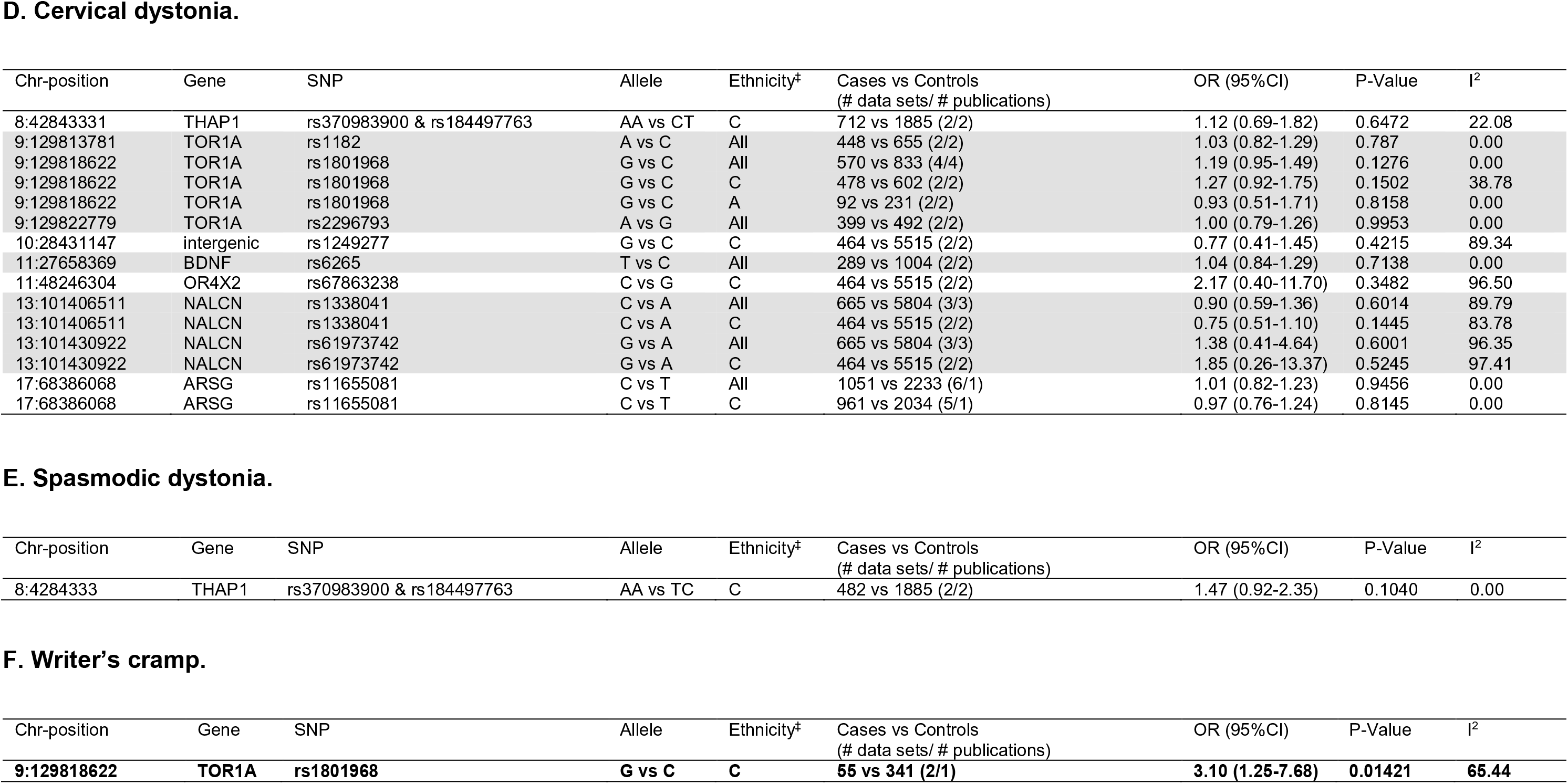
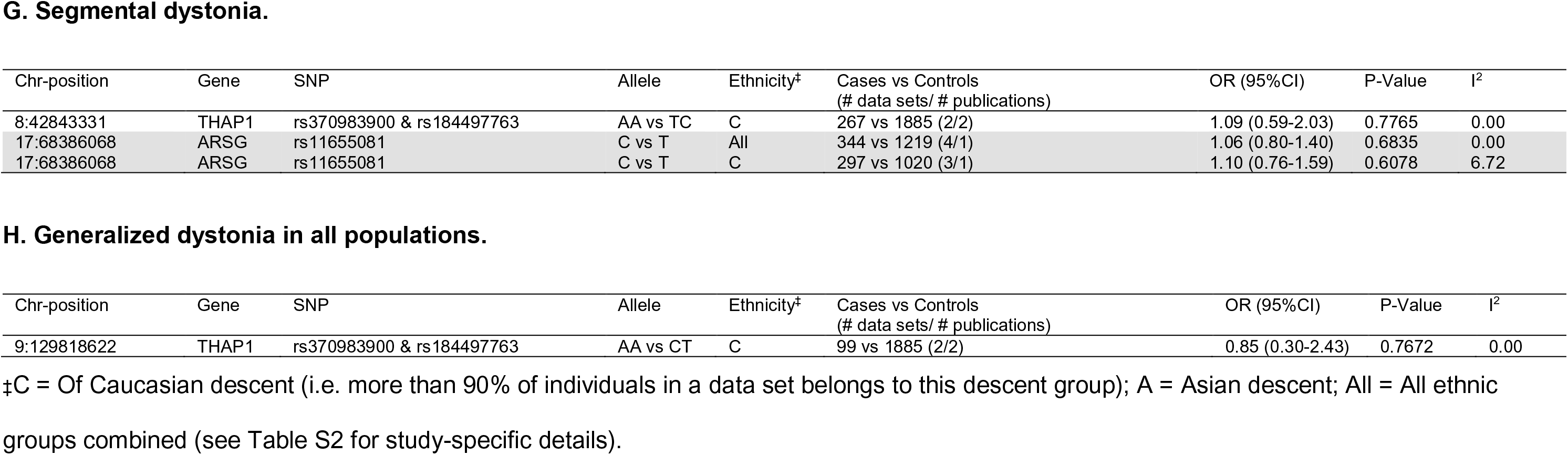
Results of random-effects meta-analyses based on an additive model in diagnostic subgroups of isolated dystonia: a) all types of focal dystonia, b) musician’s dystonia, c) blepharospasm, d) cervical dystonia, e) spasmodic dystonia, f) writer’s dystonia, g) segmental dystonia, h) generalized dystonia.

## Discussion

This work represents the first systematic field synopsis of genetic association studies in dystonia. Overall, we present the results of 134 meta-analyses on 45 variants across 17 genes. While nominal associations with several loci were observed, only variants in *DRD1* and *ARSG* survived multiple testing correction. However, even these two most significant results should be interpreted with caution owing to comparatively small sample size.

Prior to this study meta-analyses on genetic association data in dystonia were only available for two genes, i.e. *TOR1A*[9,10] and *BDNF*[11,12], both with conflicting results, i.e. one study interpreted their results in favor of a role in dystonia susceptibility, while the other study reached an opposite conclusion. Although our results provide some degree of support for an involvement of *TOR1A* SNP rs35153737 in Asians and rs13283584 in Caucasians (Table 1), no association evidence was observed with the only available (but widely tested) SNP (rs6265) in *BDNF*.

One notable observation and important limitation of our study is that most meta-analyses are based on relatively small sample sizes (median combined sample size of across all metaanalyses was only 2,882 individuals, compared to ~4,500 for Alzheimer’s[13] and ~7,500 for Parkinson’s disease[14]). Provided that the genetic liability of dystonia is mostly governed by small effect variants with ORs <1.2, our power to detect such associations was small, and only efforts with substantially increased sample sizes for both patients and controls will have sufficient power to detect such effect sizes[14]. Other limitations of our study relate to the search strategy, which might not have identified some eligible papers, and an erroneous data extraction and representation procedure. However, while such errors are unavoidable, our ample experience in the collection, annotation and curation of genetic association data in the context of systematic field synopses[13,14,24–27] suggest that these errors will likely be infrequent and will not affect the major conclusion(s) of our paper.

Notwithstanding these and possibly other unrecognized limitations, our study represents the first systematic synopsis of genetic association studies in dystonia and the largest collection of meta-analyses in the field. Despite its comprehensiveness, and in contrast to genetic studies on monogenic forms of the disease, it currently provides only little support for an existence of strong genetic risk factors acting in sporadic dystonia. This situation will likely change upon completion of additional large-scale GWAS of sufficient size, which will hopefully provide new insights into the pathogenetic forces underlying the onset and progression of this debilitating disease.

## Acknowledgements

This study was supported by by the German Research Foundation (DFG) (FOR2488, main support by subproject “P6” BE2287/6-1 to LB, additional support by subprojects “P7” LI2654/2-1 to CML, “Z1” KL1134/17-1 to CK, and “P4” LO1555/9-1 to KL). CK is the recipient of a career development award from the Hermann and Lilly Schilling Foundation. The authors report no conflict of interest.

## Supplementary Material (available from the authors upon request)

**Table S1.** Overview of search algorithms on NCBI’s “PubMed” database. In a first stage we searched for general keywords “(dystoni* OR (writer* AND cramp) OR graphospasm OR blepharospasm OR torticollis OR meige OR spasmodic OR dysphonia OR retrocollis OR antecollis OR laterocollis OR (musician* AND cramp) OR (occupation* AND cramp) OR (golf* AND cramp) OR yips) AND associat* AND gene*”. This resulted in 896 publications. In the second stage, we separately searched for each of the genes identified in stage 1 (n=10) as well as all for 28 loci with a “DYT” label (supplement table 2; DYT designations taken from refs. 5 and 15). Each of these genes was searched for the different forms of isolated dystonia “(dystoni* OR (writer* AND cramp) OR graphospasm OR blepharospasm OR torticollis OR meige OR spasmodic OR dysphonia OR retrocollis OR antecollis OR laterocollis OR (musician* AND cramp) OR (occupation* AND cramp) OR (golf* AND cramp) OR yips)” in combination with their current official HGNC (HUGO Gene Nomenclature Committee at the European Bioinformatics Institute [28] (URL: https://www.genenames.org/) gene symbol as well as all previous symbols. ¶Some authors assign *(COL6A3)* as DYT2; here, this gene is listed under DYT27.

**Table S2.** Detailed summary of genetic association studies identified for this field synopsis and meta-analyses. Tabs correspond to diagnostic subgroups of dystonia; these were also utilized for the meta-analyses.

**Table S3.** Estimates of pairwise linkage disequilibrium between meta-analyzed SNPs within the same 1Mb interval, this only related to variants in *PRKRA, DRD1, THAP1, TOR1A, DBH, DRD2, NACL, GCH1, ARSG* and *COMT* based on genotype data from the 1000 genomes project, LD was calculated using PLINK v1.9. SNP pairs with r^2^ > 0.3 were considered not independent. Bold font indicates r^2^ > 0.3.

## References

[1] H.A. Jinnah, J.K. Teller, W.R. Galpern, Recent Developments in Dystonia, (n.d.). doi:10.1097/WCO.0000000000000213.

[2] C. Klein, Genetics in dystonia, Parkinsonism Relat. Disord. 20 (2014) S137–S142. doi:10.1016/S1353-8020(13)70033-6.

[3] K. Lohmann, C. Klein, Update on the Genetics of Dystonia, (1910). doi:10.1007/s11910-017-0735-0.

[4] C. Klein, K. Lohmann, C. Marras, A. Münchau, Hereditary Dystonia Overview, University of Washington, Seattle, 1993. http://www.ncbi.nlm.nih.gov/pubmed/20301334 (accessed February 15, 2018).

[5] C. Marras, A. Lang, B.P. van de Warrenburg, C.M. Sue, S.J. Tabrizi, L. Bertram, S. Mercimek-Mahmutoglu, D. Ebrahimi-Fakhari, T.T. Warner, A. Durr, B. Assmann, K. Lohmann, V. Kostic, C. Klein, Nomenclature of genetic movement disorders: Recommendations of the international Parkinson and movement disorder society task force, Mov. Disord. 31 (2016) 436–457. doi:10.1002/mds.26527.

[6] K. Lohmann, A. Schmidt, A. Schillert, S. Winkler, A. Albanese, F. Baas, A.R. Bentivoglio, F. Borngräber, N. Brüggemann, G. Defazio, F. Del Sorbo, G. Deuschl, M.J. Edwards, T. Gasser, P. Gómez-Garre, J. Graf, J.L. Groen, A. Grünewald, J. Hagenah, C. Hemmelmann, H.C. Jabusch, R. Kaji, M. Kasten, H. Kawakami, V.S. Kostic, M. Liguori, P. Mir, A. Münchau, F. Ricchiuti, S. Schreiber, K. Siegesmund, M. Svetel, M.A.J. Tijssen, E.M. Valente, A. Westenberger, K.E. Zeuner, S. Zittel, E. Altenmüller, A. Ziegler, C. Klein, Genome-wide association study in musician’s dystonia: A risk variant at the arylsulfatase G locus?, Mov. Disord. 29 (2014) 921–927. doi:10.1002/mds.25791.

[7] K.Y. Mok, S.A. Schneider, D. Trabzuni, M. Stamelou, M. Edwards, D. Kasperaviciute, S. Pickering-Brown, M. Silverdale, J. Hardy, K.P. Bhatia, Genomewide association study in cervical dystonia demonstrates possible association with sodium leak channel., Mov. Disord. 29 (2014) 245–51. doi:10.1002/mds.25732.

[8] K. Lohmann, C. Klein, Genetics of dystonia: What’s known? What’s new? What’s next?, Mov. Disord. 28 (2013) 899–905. doi:10.1002/mds.25536.

[9] J.L. Groen, K. Ritz, M.W. Tanck, B.P. van de Warrenburg, J.J. van Hilten, M. Aramideh, F. Baas, M.A.J. Tijssen, Is TOR1A a risk factor in adult-onset primary torsion dystonia?, Mov. Disord. 28 (2013) 827–31. doi:10.1002/mds.25381.

[10] V. Siokas, E. Dardiotis, E.E. Tsironi, G. Tsivgoulis, D. Rikos, M. Sokratous, S. Koutsias, K. Paterakis, G. Deretzi, G.M. Hadjigeorgiou, The Role of TOR1A Polymorphisms in Dystonia: A Systematic Review and Meta-Analysis., PLoS One. 12 (2017) e0169934. doi:10.1371/journal.pone.0169934.

[11] P. Gómez-Garre, I. Huertas-Fernández, M.T. Cáceres-Redondo, A. Alonso-Canovas, I. Bernal-Bernal, A. Blanco-Ollero, M. Bonilla-Toribio, J.A. Burguera, M. Carballo, F. Carrillo, M.J. Catalán-Alonso, F. Escamilla-Sevilla, R. Espinosa-Rosso, M.C. Fernández-Moreno, J. García-Caldentey, J.M. García-Moreno, P.J. García-Ruiz, S. Giacometti-Silveira, J. Gutiérrez-García, S. Jesús, E. López-Valdés, J.C. Martínez-Castrillo, I. Martínez-Torres, M.P. Medialdea-Natera, C. Méndez-Lucena, A. Mínguez-Castellanos, M. Moya, J.J. Ochoa-Sepulveda, T. Ojea, N. Rodríguez, M. Sillero-Sánchez, L. Vargas-González, P. Mir, BDNF Val66Met polymorphism in primary adult-onset dystonia: A case-control study and meta-analysis, Mov. Disord. 29 (2014) 1083–1086. doi:10.1002/mds.25938.

[12] W. Sako, N. Murakami, Y. Izumi, R. Kaji, Val66Met polymorphism of brain-derived neurotrophic factor is associated with idiopathic dystonia., J. Clin. Neurosci. 22 (2015) 575–7. doi:10.1016/j.jocn.2014.08.014.

[13] L. Bertram, M.B. McQueen, K. Mullin, D. Blacker, R.E. Tanzi, Systematic metaanalyses of Alzheimer disease genetic association studies: the AlzGene database, Nat. Genet. 39 (2007) 17–23. doi:10.1038/ng1934.

[14] C.M. Lill, J.T. Roehr, M.B. McQueen, F.K. Kavvoura, S. Bagade, B.-M.M. Schjeide, L.M. Schjeide, E. Meissner, U. Zauft, N.C. Allen, T. Liu, M. Schilling, K.J. Anderson, G. Beecham, D. Berg, J.M. Biernacka, A. Brice, A.L. DeStefano, C.B. Do, N. Eriksson, S.A. Factor, M.J. Farrer, T. Foroud, T. Gasser, T. Hamza, J.A. Hardy, P. Heutink, E.M. Hill-Burns, C. Klein, J.C. Latourelle, D.M. Maraganore, E.R. Martin, M. Martinez, R.H. Myers, M.A. Nalls, N. Pankratz, H. Payami, W. Satake, W.K. Scott, M. Sharma, A.B. Singleton, K. Stefansson, T. Toda, J.Y. Tung, J. Vance, N.W. Wood, C.P. Zabetian, P. 23andMe Genetic Epidemiology of Parkinson’s Disease Consortium, R.E. International Parkinson’s Disease Genomics Consortium, M.J. Parkinson’s Disease GWAS Consortium, F. Wellcome Trust Case Control Consortium 2), P. Young, R.E. Tanzi, M.J. Khoury, F. Zipp, H. Lehrach, J.P.A. Ioannidis, L. Bertram, Comprehensive research synopsis and systematic meta-analyses in Parkinson’s disease genetics: The PDGene database., PLoS Genet. 8 (2012) e1002548. doi:10.1371/journal.pgen.1002548.

[15] E. Meyer, K.J. Carss, J. Rankin, J.M.E. Nichols, D. Grozeva, A.P. Joseph, N.E. Mencacci, A. Papandreou, J. Ng, S. Barral, A. Ngoh, H. Ben-Pazi, M.A. Willemsen, D. Arkadir, A. Barnicoat, H. Bergman, S. Bhate, A. Boys, N. Darin, N. Foulds, N. Gutowski, A. Hills, H. Houlden, J.A. Hurst, Z. Israel, M. Kaminska, P. Limousin, D. Lumsden, S. McKee, S. Misra, S.S. Mohammed, V. Nakou, J. Nicolai, M. Nilsson, H. Pall, K.J. Peall, G.B. Peters, P. Prabhakar, M.S. Reuter, P. Rump, R. Segel, M. Sinnema, M. Smith, P. Turnpenny, S.M. White, D. Wieczorek, S. Wiethoff, B.T. Wilson, G. Winter, C. Wragg, S. Pope, S.J.H. Heales, D. Morrogh, A. Pittman, L.J. Carr, B. Perez-Dueñas, J.-P. Lin, A. Reis, W.A. Gahl, C. Toro, K.P. Bhatia, N.W. Wood, E.-J. Kamsteeg, W.K. Chong, P. Gissen, M. Topf, R.C. Dale, J.R. Chubb, F.L. Raymond, M.A. Kurian, Mutations in the histone methyltransferase gene KMT2B cause complex early-onset dystonia, Nat. Genet. 49 (2017) 223–237.

[16] A. Albanese, K. Bhatia, S.B. Bressman, M.R. Delong, S. Fahn, V.S.C. Fung, M. Hallett, J. Jankovic, H.A. Jinnah, C. Klein, A.E. Lang, J.W. Mink, J.K. Teller, Phenomenology and classification of dystonia: a consensus update., Mov. Disord. 28 (2013) 863–73. doi:10.1002/mds.25475.

[17] A. Auton, G.R. Abecasis, D.M. Altshuler, R.M. Durbin, G.R. Abecasis, D.R. Bentley, A. Chakravarti, A.G. Clark, P. Donnelly, E.E. Eichler, P. Flicek, S.B. Gabriel, R.A. Gibbs, E.D. Green, M.E. Hurles, B.M. Knoppers, J.O. Korbel, E.S. Lander, C. Lee, H. Lehrach, E.R. Mardis, G.T. Marth, G.A. McVean, D.A. Nickerson, J.P. Schmidt, S.T. Sherry, J. Wang, R.K. Wilson, R.A. Gibbs, E. Boerwinkle, H. Doddapaneni, Y. Han, V. Korchina, C. Kovar, S. Lee, D. Muzny, J.G. Reid, Y. Zhu, J. Wang, Y. Chang, Q. Feng, X. Fang, X. Guo, M. Jian, H. Jiang, X. Jin, T. Lan, G. Li, J. Li, Y. Li, S. Liu, X. Liu, Y. Lu, X. Ma, M. Tang, B. Wang, G. Wang, H. Wu, R. Wu, X. Xu, Y. Yin, D. Zhang, W. Zhang, J. Zhao, M. Zhao, X. Zheng, E.S. Lander, D.M. Altshuler, S.B. Gabriel, N. Gupta, N. Gharani, L.H. Toji, N.P. Gerry, A.M. Resch, P. Flicek, J. Barker, L. Clarke, L. Gil, S.E. Hunt, G. Kelman, E. Kulesha, R. Leinonen, W.M. McLaren, R. Radhakrishnan, A. Roa, D. Smirnov, R.E. Smith, I. Streeter, A. Thormann, I. Toneva, B. Vaughan, X. Zheng-Bradley, D.R. Bentley, R. Grocock, S. Humphray, T. James, Z. Kingsbury, H. Lehrach, R. Sudbrak, M.W. Albrecht, V.S. Amstislavskiy, T.A. Borodina, M. Lienhard, F. Mertes, M. Sultan, B. Timmermann, M.-L. Yaspo, E.R. Mardis, R.K. Wilson, L. Fulton, R. Fulton, S.T. Sherry, V. Ananiev, Z. Belaia, D. Beloslyudtsev, N. Bouk, C. Chen, D. Church, R. Cohen, C. Cook, J. Garner, T. Hefferon, M. Kimelman, C. Liu, J. Lopez, P. Meric, C. O’Sullivan, Y. Ostapchuk, L. Phan, S. Ponomarov, V. Schneider, E. Shekhtman, K. Sirotkin, D. Slotta, H. Zhang, G.A. McVean, R.M. Durbin, S. Balasubramaniam, J. Burton, P. Danecek, T.M. Keane, A. Kolb-Kokocinski, S. McCarthy, J. Stalker, M. Quail, J.P. Schmidt, C.J. Davies, J. Gollub, T. Webster, B. Wong, Y. Zhan, A. Auton, C.L. Campbell, Y. Kong, A. Marcketta, R.A. Gibbs, F. Yu, L. Antunes, M. Bainbridge, D. Muzny, A. Sabo, Z. Huang, J. Wang, L.J.M. Coin, L. Fang, X. Guo, X. Jin, G. Li, Q. Li, Y. Li, Z. Li, H. Lin, B. Liu, R. Luo, H. Shao, Y. Xie, C. Ye, C. Yu, F. Zhang, H. Zheng, H. Zhu, C. Alkan, E. Dal, F. Kahveci, G.T. Marth, E.P. Garrison, D. Kural, W.-P. Lee, W. Fung Leong, M. Stromberg, A.N. Ward, J. Wu, M. Zhang, M.J. Daly, M.A. DePristo, R.E. Handsaker, D.M. Altshuler, E. Banks, G. Bhatia, G. del Angel, S.B. Gabriel, G. Genovese, N. Gupta, H. Li, S. Kashin, E.S. Lander, S.A. McCarroll, J.C. Nemesh, R.E. Poplin, S.C. Yoon, J. Lihm, V. Makarov, A.G. Clark, S. Gottipati, A. Keinan, J.L. Rodriguez-Flores, J.O. Korbel, T. Rausch, M.H. Fritz, A.M. Stütz, P. Flicek, K. Beal, L. Clarke, A. Datta, J. Herrero, W.M. McLaren, G.R.S. Ritchie, R.E. Smith, D. Zerbino, X. Zheng-Bradley, P.C. Sabeti, I. Shlyakhter, S.F. Schaffner, J. Vitti, D.N. Cooper, E. V. Ball, P.D. Stenson, D.R. Bentley, B. Barnes, M. Bauer, R. Keira Cheetham, A. Cox, M. Eberle, S. Humphray, S. Kahn, L. Murray, J. Peden, R. Shaw, E.E. Kenny, M.A. Batzer, M.K. Konkel, J.A. Walker, D.G. MacArthur, M. Lek, R. Sudbrak, V.S. Amstislavskiy, R. Herwig, E.R. Mardis, L. Ding, D.C. Koboldt, D. Larson, K. Ye, S. Gravel, A. Swaroop, E. Chew, T. Lappalainen, Y. Erlich, M. Gymrek, T. Frederick Willems, J.T. Simpson, M.D. Shriver, J.A. Rosenfeld, C.D. Bustamante, S.B. Montgomery, F.M. De La Vega, J.K. Byrnes, A.W. Carroll, M.K. DeGorter, P. Lacroute, B.K. Maples, A.R. Martin, A. Moreno-Estrada, S.S. Shringarpure, F. Zakharia, E. Halperin, Y. Baran, C. Lee, E. Cerveira, J. Hwang, A. Malhotra, D. Plewczynski, K. Radew, M. Romanovitch, C. Zhang, F.C.L. Hyland, D.W. Craig, A. Christoforides, N. Homer, T. Izatt, A.A. Kurdoglu, S.A. Sinari, K. Squire, S.T. Sherry, C. Xiao, J. Sebat, D. Antaki, M. Gujral, A. Noor, K. Ye, E.G. Burchard, R.D. Hernandez, C.R. Gignoux, D. Haussler, S.J. Katzman, W. James Kent, B. Howie, A. Ruiz-Linares, E.T. Dermitzakis, S.E. Devine, G.R. Abecasis, H. Min Kang, J.M. Kidd, T. Blackwell, S. Caron, W. Chen, S. Emery, L. Fritsche, C. Fuchsberger, G. Jun, B. Li, R. Lyons, C. Scheller, C. Sidore, S. Song, E. Sliwerska, D. Taliun, A. Tan, R. Welch, M. Kate Wing, X. Zhan, P. Awadalla, A. Hodgkinson, Y. Li, X. Shi, A. Quitadamo, G. Lunter, G.A. McVean, J.L. Marchini, S. Myers, C. Churchhouse, O. Delaneau, A. Gupta-Hinch, W. Kretzschmar, Z. Iqbal, I. Mathieson, A. Menelaou, A. Rimmer, D.K. Xifara, T.K. Oleksyk, Y. Fu, X. Liu, M. Xiong, L. Jorde, D. Witherspoon, J. Xing, E.E. Eichler, B.L. Browning, S.R. Browning, F. Hormozdiari, P.H. Sudmant, E. Khurana, R.M. Durbin, M.E. Hurles, C. Tyler-Smith, C.A. Albers, Q. Ayub, S. Balasubramaniam, Y. Chen, V. Colonna, P. Danecek, L. Jostins, T.M. Keane, S. McCarthy, K. Walter, Y. Xue, M.B. Gerstein, A. Abyzov, S. Balasubramanian, J. Chen, D. Clarke, Y. Fu, A.O. Harmanci, M. Jin, D. Lee, J. Liu, X. Jasmine Mu, J. Zhang, Y. Zhang, Y. Li, R. Luo, H. Zhu, C. Alkan, E. Dal, F. Kahveci, G.T. Marth, E.P. Garrison, D. Kural, W.-P. Lee, A.N. Ward, J. Wu, M. Zhang, S.A. McCarroll, R.E. Handsaker, D.M. Altshuler, E. Banks, G. del Angel, G. Genovese, C. Hartl, H. Li, S. Kashin, J.C. Nemesh, K. Shakir, S.C. Yoon, J. Lihm, V. Makarov, J. Degenhardt, J.O. Korbel, M.H. Fritz, S. Meiers, B. Raeder, T. Rausch, A.M. Stütz, P. Flicek, F. Paolo Casale, L. Clarke, R.E. Smith, O. Stegle, X. Zheng-Bradley, D.R. Bentley, B. Barnes, R. Keira Cheetham, M. Eberle, S. Humphray, S. Kahn, L. Murray, R. Shaw, E.-W. Lameijer, M.A. Batzer, M.K. Konkel, J.A. Walker, L. Ding, I. Hall, K. Ye, P. Lacroute, C. Lee, E. Cerveira, A. Malhotra, J. Hwang, D. Plewczynski, K. Radew, M. Romanovitch, C. Zhang, D.W. Craig, N. Homer, D. Church, C. Xiao, J. Sebat, D. Antaki, V. Bafna, J. Michaelson, K. Ye, S.E. Devine, E.J. Gardner, G.R. Abecasis, J.M. Kidd, R.E. Mills, G. Dayama, S. Emery, G. Jun, X. Shi, A. Quitadamo, G. Lunter, G.A. McVean, K. Chen, X. Fan, Z. Chong, T. Chen, D. Witherspoon, J. Xing, E.E. Eichler, M.J. Chaisson, F. Hormozdiari, J. Huddleston, M. Malig, B.J. Nelson, P.H. Sudmant, N.F. Parrish, E. Khurana, M.E. Hurles, B. Blackburne, S.J. Lindsay, Z. Ning, K. Walter, Y. Zhang, M.B. Gerstein, A. Abyzov, J. Chen, D. Clarke, H. Lam, X. Jasmine Mu, C. Sisu, J. Zhang, Y. Zhang, R.A. Gibbs, F. Yu, M. Bainbridge, D. Challis, U.S. Evani, C. Kovar, J. Lu, D. Muzny, U. Nagaswamy, J.G. Reid, A. Sabo, J. Yu, X. Guo, W. Li, Y. Li, R. Wu, G.T. Marth, E.P. Garrison, W. Fung Leong, A.N. Ward, G. del Angel, M.A. DePristo, S.B. Gabriel, N. Gupta, C. Hartl, R.E. Poplin, A.G. Clark, J.L. Rodriguez-Flores, P. Flicek, L. Clarke, R.E. Smith, X. Zheng-Bradley, D.G. MacArthur, E.R. Mardis, R. Fulton, D.C. Koboldt, S. Gravel, C.D. Bustamante, D.W. Craig, A. Christoforides, N. Homer, T. Izatt, S.T. Sherry, C. Xiao, E. T. Dermitzakis, G.R. Abecasis, H. Min Kang, G.A. McVean, M.B. Gerstein, S. Balasubramanian, L. Habegger, H. Yu, P. Flicek, L. Clarke, F. Cunningham, I. Dunham, D. Zerbino, X. Zheng-Bradley, K. Lage, J. Berg Jespersen, H. Horn, S.B. Montgomery, M.K. DeGorter, E. Khurana, C. Tyler-Smith, Y. Chen, V. Colonna, Y. Xue, M.B. Gerstein, S. Balasubramanian, Y. Fu, D. Kim, A. Auton, A. Marcketta, R. Desalle, A. Narechania, M.A. Wilson Sayres, E.P. Garrison, R.E. Handsaker, S. Kashin, S.A. McCarroll, J.L. Rodriguez-Flores, P. Flicek, L. Clarke, X. Zheng-Bradley, Y. Erlich, M. Gymrek, T. Frederick Willems, C.D. Bustamante, F.L. Mendez, G. David Poznik, P.A. Underhill, C. Lee, E. Cerveira, A. Malhotra, M. Romanovitch, C. Zhang, G.R. Abecasis, L. Coin, H. Shao, D. Mittelman, C. Tyler-Smith, Q. Ayub, R. Banerjee, M. Cerezo, Y. Chen, T.W. Fitzgerald, S. Louzada, A. Massaia, S. McCarthy, G.R. Ritchie, Y. Xue, F. Yang, R.A. Gibbs, C. Kovar, D. Kalra, W. Hale, D. Muzny, J.G. Reid, J. Wang, X. Dan, X. Guo, G. Li, Y. Li, C. Ye, X. Zheng, D.M. Altshuler, P. Flicek, L. Clarke, X. Zheng-Bradley, D.R. Bentley, A. Cox, S. Humphray, S. Kahn, R. Sudbrak, M.W. Albrecht, M. Lienhard, D. Larson, D.W. Craig, T. Izatt, A.A. Kurdoglu, S.T. Sherry, C. Xiao, D. Haussler, G.R. Abecasis, G.A. McVean, R.M. Durbin, S. Balasubramaniam, T.M. Keane, S. McCarthy, J. Stalker, A. Chakravarti, B.M. Knoppers, G.R. Abecasis, K.C. Barnes, C. Beiswanger, E.G. Burchard, C.D. Bustamante, H. Cai, H. Cao, R.M. Durbin, N.P. Gerry, N. Gharani, R.A. Gibbs, C.R. Gignoux, S. Gravel, B. Henn, D. Jones, L. Jorde, J.S. Kaye, A. Keinan, A. Kent, A. Kerasidou, Y. Li, R. Mathias, G.A. McVean, A. Moreno-Estrada, P.N. Ossorio, M. Parker, A.M. Resch, C.N. Rotimi, C.D. Royal, K. Sandoval, Y. Su, R. Sudbrak, Z. Tian, S. Tishkoff, L.H. Toji, C. Tyler-Smith, M. Via, Y. Wang, H. Yang, L. Yang, J. Zhu, W. Bodmer, G. Bedoya, A. Ruiz-Linares, Z. Cai, Y. Gao, J. Chu, L. Peltonen, A. Garcia-Montero, A. Orfao, J. Dutil, J.C. Martinez-Cruzado, T.K. Oleksyk, K.C. Barnes, R.A. Mathias, A. Hennis, H. Watson, C. McKenzie, F. Qadri, R. LaRocque, P.C. Sabeti, J. Zhu, X. Deng, P.C. Sabeti, D. Asogun, O. Folarin, C. Happi, O. Omoniwa, M. Stremlau, R. Tariyal, M. Jallow, F. Sisay Joof, T. Corrah, K. Rockett, D. Kwiatkowski, J. Kooner, T. Tinh Hiê’n, S.J. Dunstan, N. Thuy Hang, R. Fonnie, R. Garry, L. Kanneh, L. Moses, P.C. Sabeti, J. Schieffelin, D.S. Grant, C. Gallo, G. Poletti, D. Saleheen, A. Rasheed, L.D. Brooks, A.L. Felsenfeld, J.E. McEwen, Y. Vaydylevich, E.D. Green, A. Duncanson, M. Dunn, J.A. Schloss, J. Wang, H. Yang, A. Auton, L.D. Brooks, R.M. Durbin, E.P. Garrison, H. Min Kang, J.O. Korbel, J.L. Marchini, S. McCarthy, G.A. McVean, G.R. Abecasis, A global reference for human genetic variation, Nature. 526 (2015) 68–74. doi:10.1038/nature15393.

[18] J.R.B. Newman, G.T. Sutherland, R.S. Boyle, N. Limberg, S. Blum, J.D. O’Sullivan, P.A. Silburn, G.D. Mellick, Common polymorphisms in dystonia-linked genes and susceptibility to the sporadic primary dystonias., Parkinsonism Relat. Disord. 18 (2012) 351–7. doi:10.1016/j.parkreldis.2011.11.024.

[19] A. Djarmati, S.A. Schneider, K. Lohmann, S. Winkler, H. Pawlack, J. Hagenah, N. Brüggemann, S. Zittel, T. Fuchs, A. Raković, A. Schmidt, H.-C. Jabusch, R. Wilcox, V.S. Kostić, H. Siebner, E. Altenmüller, A. Münchau, L.J. Ozelius, C. Klein, Mutations in THAP1 (DYT6) and generalised dystonia with prominent spasmodic dysphonia: a genetic screening study, Lancet Neurol. 8 (2009) 447–452. doi:10.1016/S1474-4422(09)70083-3.

[20] J.R.B. Newman, M. Todorovic, P.A. Silburn, G.T. Sutherland, G.D. Mellick, Lack of reproducibility in re-evaluating associations between GCH1 polymorphisms and Parkinson’s disease and isolated dystonia in an Australian case-control group, Parkinsonism Relat. Disord. 20 (2014) 668–670. doi:10.1016/j.parkreldis.2014.02.014.

[21] K. Lohmann, N. Uflacker, A. Erogullari, T. Lohnau, S. Winkler, A. Dendorfer, S.A. Schneider, A. Osmanovic, M. Svetel, A. Ferbert, S. Zittel, A.A. Kühn, A. Schmidt, E. Altenmüller, A. Münchau, C. Kamm, M. Wittstock, A. Kupsch, E. Moro, J. Volkmann, V. Kostic, F.J. Kaiser, C. Klein, N. Brüggemann, Identification and functional analysis of novel THAP1 mutations., Eur. J. Hum. Genet. 20 (2012) 171–5.

[22] S. Purcell, B. Neale, K. Todd-Brown, L. Thomas, M.A.R. Ferreira, D. Bender, J. Maller, P. Sklar, P.I.W. de Bakker, M.J. Daly, P.C. Sham, PLINK: a tool set for whole-genome association and population-based linkage analyses., Am. J. Hum. Genet. 81 (2007) 559–75. doi:10.1086/519795.

[23] C.C. Chang, C.C. Chow, L.C. Tellier, S. Vattikuti, S.M. Purcell, J.J. Lee, Second-generation PLINK: rising to the challenge of larger and richer datasets, Gigascience. 4 (2015) 7. doi:10.1186/s13742-015-0047-8.

[24] N.C. Allen, S. Bagade, M.B. McQueen, J.P.A. Ioannidis, F.K. Kavvoura, M.J. Khoury, R.E. Tanzi, L. Bertram, Systematic meta-analyses and field synopsis of genetic association studies in schizophrenia: the SzGene database., Nat. Genet. 40 (2008) 827–34. doi:10.1038/ng.171.

[25] P.J. Castaldi, M.H. Cho, M. Cohn, F. Langerman, S. Moran, N. Tarragona, H. Moukhachen, R. Venugopal, D. Hasimja, E. Kao, B. Wallace, C.P. Hersh, S. Bagade, L. Bertram, E.K. Silverman, T.A. Trikalinos, The COPD genetic association compendium: a comprehensive online database of COPD genetic associations., Hum. Mol. Genet. 19 (2010) 526–34. doi:10.1093/hmg/ddp519.

[26] F. Chatzinasiou, C.M. Lill, K. Kypreou, I. Stefanaki, V. Nicolaou, G. Spyrou, E. Evangelou, J.T. Roehr, E. Kodela, A. Katsambas, H. Tsao, J.P.A. Ioannidis, L. Bertram, A.J. Stratigos, Comprehensive Field Synopsis and Systematic Meta-analyses of Genetic Association Studies in Cutaneous Melanoma, JNCI J. Natl. Cancer Inst. 103 (2011) 1227–1235. doi:10.1093/jnci/djr219.

[27] C.M. Lill, O. Abel, L. Bertram, A. Al-Chalabi, Keeping up with genetic discoveries in amyotrophic lateral sclerosis: the ALSoD and ALSGene databases., Amyotroph. Lateral Scler. 12 (2011) 238–49. doi:10.3109/17482968.2011.584629.

[28] B. Yates, B. Braschi, K.A. Gray, R.L. Seal, S. Tweedie, E.A. Bruford, Genenames.org: the HGNC and VGNC resources in 2017., Nucleic Acids Res. 45 (2017) D619–D625. doi:10.1093/nar/gkw1033.

